# Hunting Eagles with glass mice: revisiting the inoculum effect for *Streptococcus pyogenes* with a hollow fibre infection model

**DOI:** 10.1101/2021.01.07.425826

**Authors:** Darcy Marum, Laurens Manning, Edward Raby

## Abstract

**Background:** Severe, invasive *Streptococcus pyogenes* (Strep A) infections result in greater than 500,000 deaths annually. First line treatment for such infections is combination benzylpenicillin and clindamycin, but treatment failure can occur with this regimen. This failure has been partially attributed to the inoculum effect, which presents as reduced antibiotic susceptibility during high bacterial density and plateau-phase growth. Hollow fibre infection models (HFIM) have been proposed as an alternative to *in vivo* research to study these effects.

**Objectives:** To re-evaluate the inoculum effect for benzylpenicillin, clindamycin, linezolid and trimethoprim-sulfamethoxazole using a Strep A HFIM.

**Methods:** Differential antibiotic susceptibility of Strep A was measured in a HFIM starting from low- and high-density inocula. Dynamic antibiotic concentrations were delivered over 48 hours to simulate human pharmacokinetics. Differences in antibiotic susceptibility were determined at 24 and 48 hours by plate count of remaining viable colony-forming units.

**Results:** Inoculum effects were seen in benzylpenicillin and linezolid at 24 hours, and benzylpenicillin, linezolid and clindamycin at 48 hours. The effect size was greatest for continuously infused benzylpenicillin. No inoculum effect was seen in trimethoprim-sulfamethoxazole.

**Conclusions:** Inoculum effects were seen in the HFIM model using benzylpenicillin, linezolid and clindamycin, which may predict reduced clinical efficacy following treatment delay. The model has proven robust and largely in agreeance with published data, recommending it for further Strep A study.

## Introduction

*Streptococcus pyogenes* (Strep A) is an important human pathogen with diverse manifestations ranging from mild pharyngitis and skin infections to life-threatening invasive disease.^1^ The burden of disease in Australia remains high, particularly among the Indigenous population, and disease mitigation efforts are hampered by inconsistent mandatory reporting requirements^2^. Penicillin remains the cornerstone antibiotic for established infections and, in the absence of an effective vaccine, for subsequent secondary prevention. However, for severe infections a second antibiotic such as clindamycin is commonly added to improve clinical outcomes, an approach often justified by the understanding that bacteria may display differential susceptibility to antibiotics at different population densities and growth phases.^3^ This phenomenon, termed the ‘inoculum effect’ was first described *in vivo* by Harry Eagle in 1952 who reported a murine myositis model of Strep A infection which showed that mice with a higher initial inoculum had decreased survival.^4^ Using the same animal infection model, and with findings replicated in static *in vitro* models, subsequent studies have demonstrated that the addition of clindamycin significantly improved outcomes. Stevens et al. demonstrated this effect was more pronounced with increasing inoculum or treatment delay.^5, 6^ In parallel, minimum inhibitory concentrations (MIC) rose with increasing inoculum.^6^ An *in vitro* inoculum effect has also been observed for clindamycin and vancomycin in the treatment of *Staphylococcus aureus* and has been proposed as a mechanistic explanation for improved outcomes observed with dual therapy with these agents for severe skin and soft tissue infections.^7, 8^

Extrapolating the results of mouse and static *in vitro* models to clinically relevant effects in humans may not necessarily be valid and should be interpreted in the context of likely *in vivo* antibiotic exposures. In the studies by Eagle and Stevens, the penicillin dosing and duration of therapy relative to the MIC did not replicate typical unbound antibiotic exposures observed in human infections treated with contemporary intravenous doses.^4, 6^

To explore the implications of this, we established a dynamic hollow fibre infection model (HFIM) of Strep A infection. This HFIM was applied to accurately replicate *in vivo* antibiotic exposure profiles typically seen with antibiotic therapy (figure 1). In this experiment, the inoculum effect was revisited using a well-characterised Strep A strain exposed to benzylpenicillin, clindamycin, linezolid, and trimethoprim/sulfamethoxazole (SXT). The latter two antibiotics were included as future alternatives for clindamycin to which resistance is rising^9^, in particular linezolid which has previously been demonstrated as effective in the treatment of necrotising Strep A infections^10^. The activity of SXT against Strep A has historically been questioned but recent *in vitro* and human data demonstrate a clinically relevant effect.^11, 12^ Given increasing interest in delivering beta-lactam therapy by continuous infusion in intensive care or outpatient settings, we also compared this route of administration with conventional bolus dosing of benzylpenicillin.

**Figure 1.**
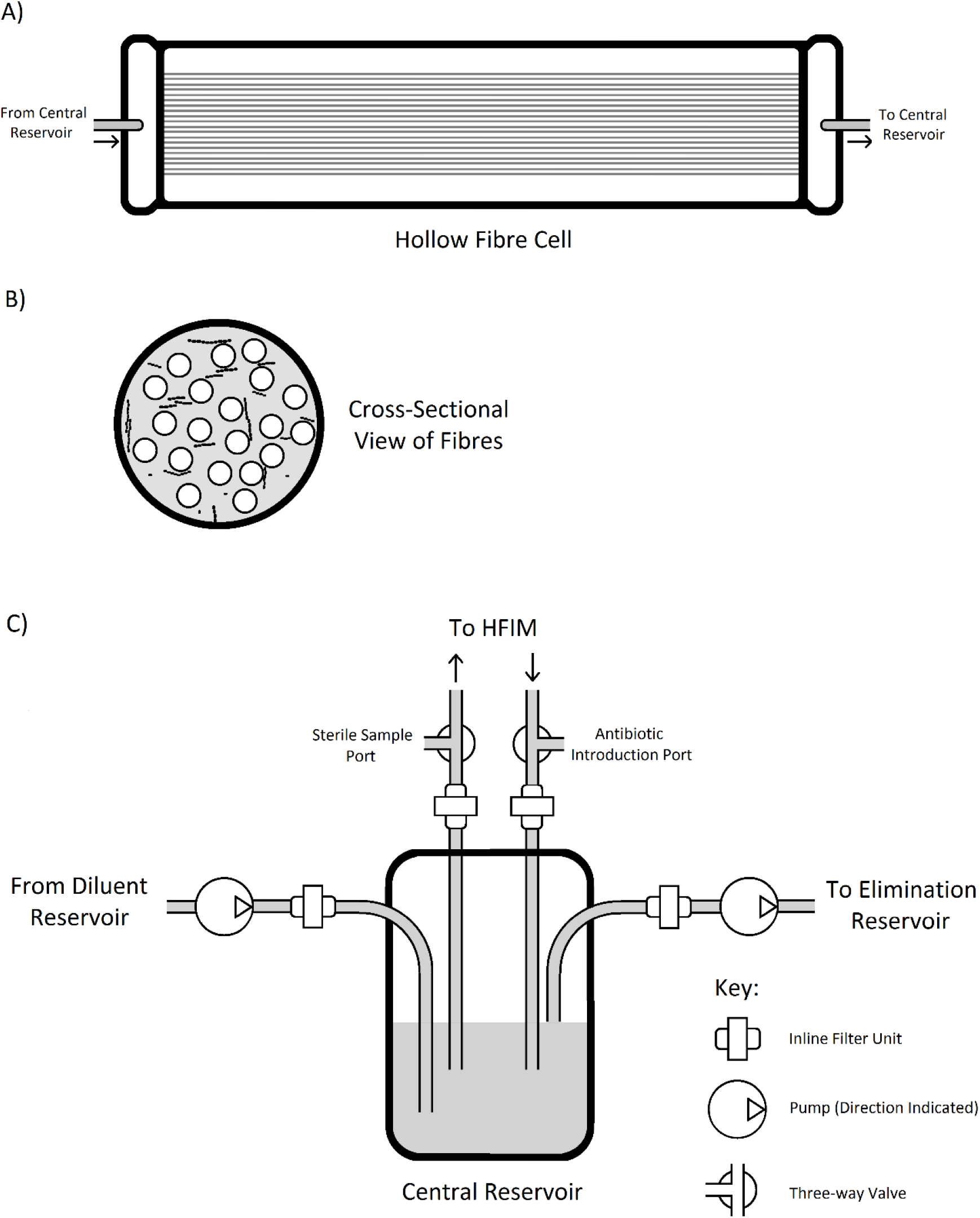
The central reservoir of the hollow fibre infection model with associated components. (A) Detail of the hollow fibre cell itself. Inputs are received on the left from the central reservoir and recirculated back to this reservoir on the right. (B) Cross-sectional view of a section of the hollow fibre cell. Bacteria are retained in the extracapillary space (grey) while media flows through the capillaries (white). (C) Outlines the overall layout of the HFIM.

## Materials and Methods

### Characterisation of Strep A isolate

Isolates were cultured and introduced to the HFIM from a frozen glycerol stock of strain HKU488 (BioSample: SAMEA1523579^*^). This isolate is an example of a globally-distributed strain expressing M-type 1, and notable as a cause of high rates of invasive and disseminated infection.^13^ Using previously described broth dilution techniques,^14^ the MICs for benzylpenicillin, clindamycin, linezolid and SXT were confirmed to be 0.005625, 0.045, 2 and 0.64 mg/L respectively, indicating susceptibility to these antibiotics as per EUCAST criteria.^15^

### Standard Growth Curve and Growth Phases in the HFIM

A thawed sample (0.05mL) was cultured on HBA from frozen glycerol stock of strain HKU488 (24 hours, 37°C, 5% CO_2_). A sample from a single colony was introduced to the HFIM running under standard experimental conditions (37°C with media recirculation via peristaltic pump). Bacterial densities were determined via dilutional plate count in triplicate at 0, 4, 8, 24, 32, 74 and 96 hours. Growth phases were determined by analysis of the resulting growth curve (figure 3).

### Pharmacokinetic parameters

Pharmacokinetic attributes were modelled to emulate *in vivo* unbound antibiotic exposure conditions expected in human dosing regimens (table 1).

**Table 1.** Predicted pharmacokinetic profiles of antibiotics in hollow fibre infection model of *Streptococcus pyogenes*

| Antibiotic | t<sub>1/2</sub><br>(Hours) | C<sub>max</sub><br>(mg/L) | Protein Bound<br>(1 - f) (%) | fC<sub>max</sub><br>(mg/L) | fC<sub>min</sub><br>(mg/L) | fAUC<sub>0,24</sub><br>(mg/L·h) | t > MIC<br>(%) | t > MBC<br>(%) | Reference |
| --- | --- | --- | --- | --- | --- | --- | --- | --- | --- |
| Benzylopenicillin<br>(Intermittent) | 0.7 | 400 | 0.60 | 160 | 3.21 | 980 | 100 | 100 | <sup>26</sup> |
| Benzylopenicillin<br>(Continuous) | - | 20 | 0.60 | 8 | 8 | 192 | 100 | 100 | <sup>26</sup> |
| Clindamycin | 4 | 8.3 | 0.77 | 1.91 | 0.48 | 10.84 | 100 | - | <sup>27</sup> |
| Linezolid | 5 | 13.4 | 0.13 | 11.66 | 2.23 | 149.50 | 100 | 0 | <sup>27</sup> |
| Trimethoprim-Sulfamethoxazole | 10 | 40 | 0.53 | 18 | 7.87 | 357.20 | 100 | - | <sup>27</sup> |

### Sample Preparation and Acclimatisation

Strep A samples were derived from a common culture of HKU488. A single colony from 48-hour growth on horse blood agar (HBA) was incubated at 37°C in CAMHB for 24 hours to reach the early plateau phase of growth. High (≥10^8^ cfu/mL) and low (≤10^7^ cfu/mL) inocula were obtained by introduction of 10mL and 100μL, respectively into each hollow fibre flow cell to produce an initial hundred-fold difference in concentration. The bacteria were then acclimatised to the HFIM under standard experimental conditions (37°C with media recirculation via peristaltic pump at 14 rpm) for 16 hours, allowing the low inoculum group to re-enter exponential phase growth while the high inoculum remained in plateau phase. Bacterial densities were measured by dilutional plate count in triplicate immediately prior to antibiotic introduction (table 2).

**Table 2.** Measured bacterial concentrations in cfu/mL immediately prior to antibiotic introduction for each antibiotic, divided into high and low initial inoculum groups.

| Antibiotic | Benzylopenicillin<br>(Continuous) |  | Benzylopenicillin<br>(Bolus) |  | Clindamycin |  | Linezolid |  | SXT |  |
| --- | --- | --- | --- | --- | --- | --- | --- | --- | --- | --- |
|  | High | Low | High | Low | High | Low | High | Low | High | Low |
| cfu/mL<br>(point estimate) | 1.02E+09 | 1.58E+07 | 4.03E+08 | 7.87E+07 | 2.75E+08 | 9.56E+06 | 8.15E+08 | 2.50E+07 | 9.99E+09 | 6.70E+07 |

Antibiotics were introduced at defined doses and intervals reflecting clinical dosing regimens (table 3) and the viable cfu/mL serially measured over 48 hours following antibiotic exposure. Samples (1mL) were collected aseptically from the flow cell at 0, 4, 8, 24, 32, and 48 hours, centrifugally pelleted (6000 RCF, 3 minutes) and resuspended in isotonic saline to preserve viability and remove residual antibiotic.

**Table 3.** Antibiotic dose and frequency together with central reservoir volume and pump settings for different antibiotics in hollow fibre model of *Streptococcus pyogenes*

| Antibiotic | Benzylpenicillin<br>(Bolus Dosed) | Benzylpenicillin<br>(Continuous) | Clindamycin | Linezolid | Trimethoprim-<br>sulphamethoxazole |
| --- | --- | --- | --- | --- | --- |
| Antibiotic Dose | 20 mg | Maintained at a free<br>concentration of 20 mg/L | 0.38 mg | 2.33 mg | 0.23 mg |
| Dose Frequency | 4 Hours | - | 8 Hours | 12 Hours | 12 Hours |
| Central Reservoir<br>Volume | 125 mL | 200 mL | 200 mL | 200 mL | 200 mL |
| Peristaltic Pump RPM | 41.25 | 10 | 11.55 | 9.24 | 4.62 |

### Viable cell density enumeration by plate count and microscopy

The density of viable bacteria following antibiotic exposure was measured from each sample. Bacterial concentration was estimated via NanoDrop 2000c (ThermoFisher Scientific) optical density readings and a defined-volume fluorescent cell counting method, and dilutional plate counts (HBA) were made in triplicate based on this estimation. Colony forming units were counted following plate incubation (24 hours, 37°C, 5% CO_2_) and expressed as log_10_ decrease in cfu/mL from baseline. From these measurements, the difference in bactericidal activity between low- and high-inoculum groups was calculated. Due to a lack of consistent definitions across different studies, we considered a difference of greater than 10 times (1 log_10_-fold) to represent an inoculum effect.^3, 16, 17^

## Results

At 24 hours, a moderate difference between high and low inocula was observed for bolus-dosed benzylpenicillin; a log_10_-fold difference of 1.34. A greater log_10_-fold difference of 3.78 was seen in the continuously infused benzylpenicillin group. Modest differences between high and low inocula were observed with clindamycin (0.79) and linezolid (1.50) but no significant difference with SXT (0.17). Notably, benzylpenicillin showed by far the greatest reduction in cfu/mL at 24 hours in the low inoculum arm but continuously infused benzylpenicillin and linezolid showed less than 1 log_10_ reduction in the high inoculum arm (table 4 and figure 2).

**Table 4.**
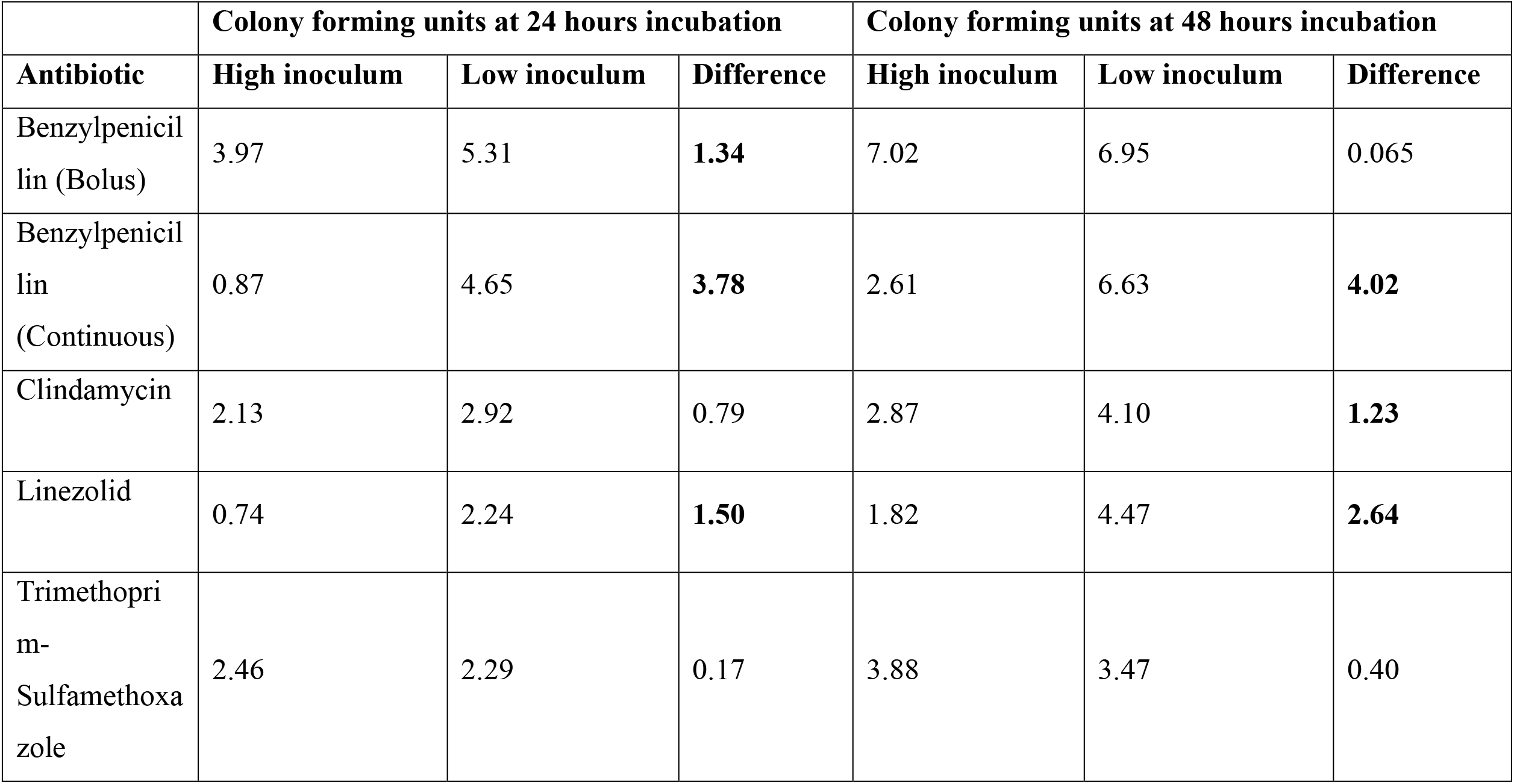
Decrease in colony forming units at high and low inocula at 24 and 48 hours following different antibiotics in the hollow fibre infection model of *Streptococcus pyogenes.* All units are expressed in terms of log_10_-fold cfu/mL decrease from initial concentration.

**Figure 2.**
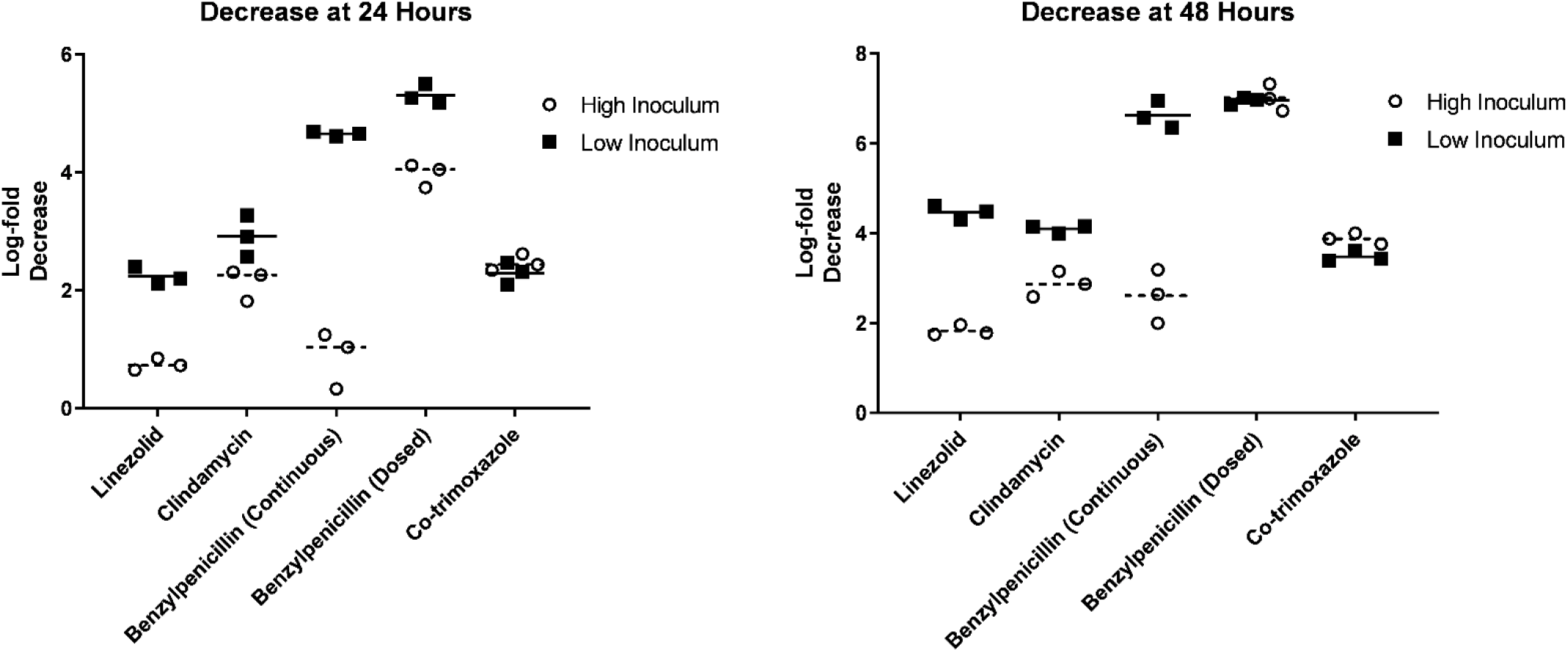
Decrease in cfu/mL between experimental groups with high and low initial inocula at 24 and 48 hours, respectively.

**Figure 3.**
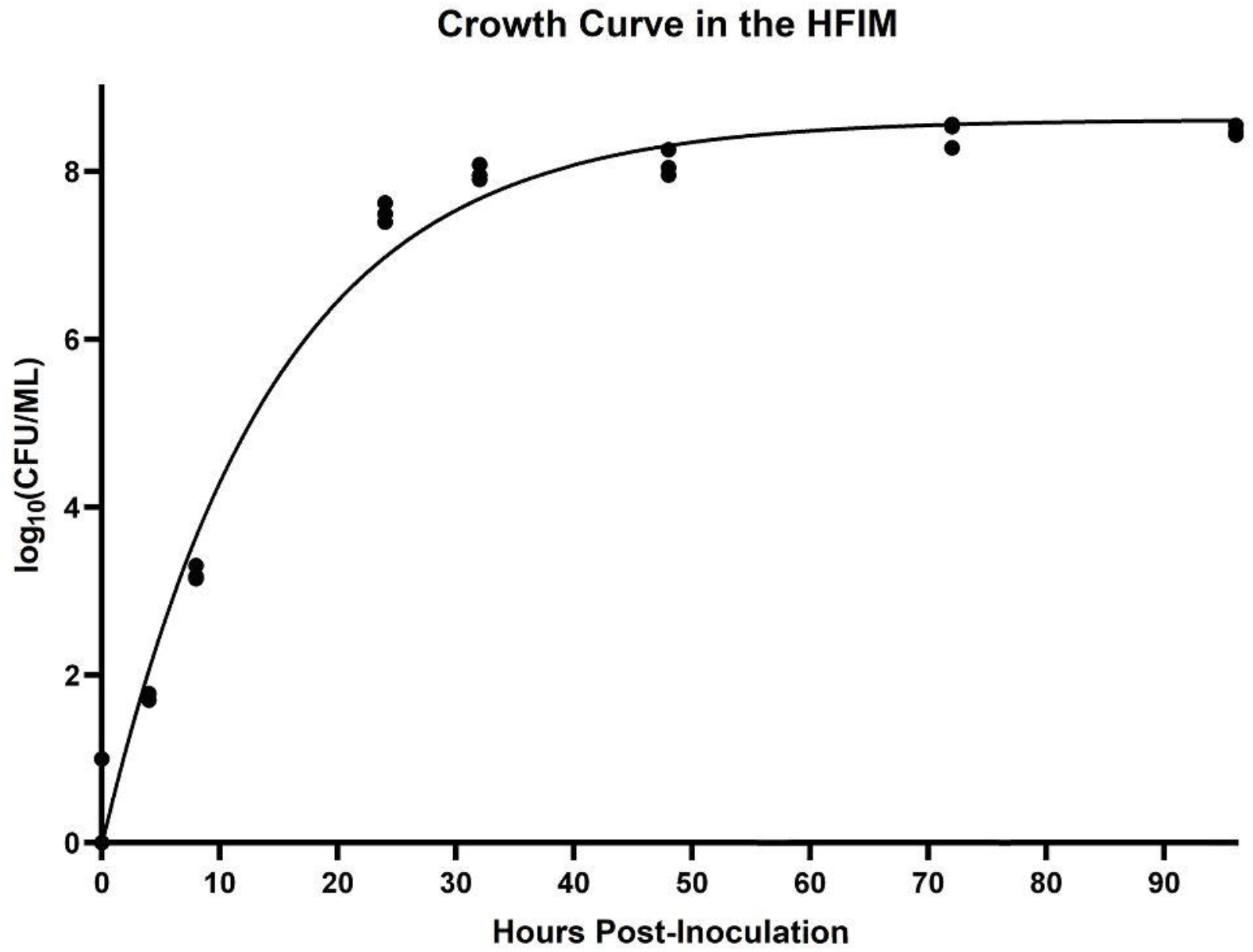
growth curve of strep A in the HFIM with no antibiotic. Regression was added via the method of least squares as a one-phase decay curve. In this model, bacteria enter the plateau phase at approximately 24 hours, at a density of 10^8^ cfu/mL.

At 48 hours there was no observed difference (0.065) between low and high inoculum arms in the bolus-dosed benzylpenicillin group in contrast to continuous-infusion benzylpenicillin which showed a log_10_-fold difference of 4.02. The pattern was otherwise similar with modest differences for clindamycin (1.23) and linezolid (2.64) and minimal difference for SXT (0.40).

## Discussion

### Definition of Inoculum Effect

Heterogeneity exists in the literature^16^ regarding how to define the inoculum effect; as such, a 10-fold change in viable bacterial concentration between groups at 24 and 48 hours was chosen as an appropriate effect size. Others have quantified this effect as either an increase in MIC,^3^ or as a time-kill rate.^17^

### Inoculum Effect of Penicillin

A large inoculum effect was observed with benzylpenicillin delivered as a continuous infusion at 24 hours, with a substantial difference in bactericidal activity between low and high inocula. This has implications for the use of penicillin (and beta-lactams generally) in the treatment of established infections, or in other cases of high bacterial load such as following delayed antibiotic administration. These results emphasise the importance of early intervention, or possibly the selection of an alternate antibiotic class unaffected by the inoculum effect, if this is not possible.

Bolus-dosed benzylpenicillin achieved high bactericidal activity in both high and low inoculum groups, only displaying an inoculum effect at 24 hours. The likely explanation is a feature of this HFIM specific to the experimental requirements of bolus-dosed benzylpenicillin. To simulate the *in vivo* clearance rate of benzylpenicillin, large volumes of CAMHB were used, approaching four litres per day (compared to less than one litre for other experiments). While necessary to simulate physiologically accurate clearance rates, this continuous provision of fresh media may have hindered both nutritional depletion and waste product accumulation that have been hypothesised as the drivers of the inoculum effect.^4^ It is possible that high flow rates also removed quorum sensing and other bacterial signalling molecules. These molecules induce changes in metabolic phenotypes associated with variation in antimicrobial resistance. In particular, the LuxS/AI-2 signalling system has been shown to alter Strep A virulence factor expression and capsule production, ^18, 19^ and is known to modulate penicillin resistance in *Streptococcus pneumoniae* and *Staphylococcus aureus.*^20, 21^

This hypothesis is supported by the observation of an inoculum effect at 24 hours. At this point in the experiment bacterial density (and the resulting metabolites, waste products and quorum sensing molecules) was much higher, and it is likely that this counteracted the high media flow. This suggests that the HFIM is best suited to high bacterial density experiments, or to those involving antibiotics with lower *in vivo* clearance rates.

### Clindamycin Inoculum Effect

Significant differences in bactericidal activity were observed at 24 and 48 hours. Current *in vivo* studies are mixed in their prediction of an inoculum effect - previous models of mouse myositis caused by Strep A infection did not show a strong correlation between inoculum size and bacterial survival,^5^ while *in vitro* studies have demonstrated^6^ an increase in MBC in higher density populations, although not necessarily an inoculum effect. It is likely that these inconsistent *in vitro* and *in vivo* results stem from clindamycin’s inhibition of bacterial exotoxins,^22^ which may modulate both bacterial virulence and the immune response *in vivo* which is not modelled in the HFIM.

### Linezolid Inoculum Effect

A significant inoculum effect was observed with linezolid at both 24 and 48 hours. While current *in vivo* studies detailing this effect are lacking in Strep A, this result accords with previous *in vivo* studies of a murine streptococcal infection model using *Streptococcus pneumoniae.*^*23*^ Bactericidal activity was no more than that achieved by benzylpenicillin against the high inoculum, suggesting that linezolid is a useful adjunct to, but not a replacement for penicillin.

### Lack of Trimethoprim/sulfamethoxazole Inoculum Effect

There was no inoculum effect demonstrated in trimethoprim/sulfamethoxazole. Inoculum effects in SXT have been previously observed in Strep A *in vitro*^*24*^, however, current *in vivo* studies are unavailable and further research is needed. If the absence of the inoculum effect is reflected in *in vivo* data, SXT may find use in the treatment of late-stage infections with high bacterial density which are not sufficiently susceptible to other antibiotics.

### Bacterial Acclimatisation and Separation of cfu/mL Values

An extended period of bacterial acclimatisation within the HFIM was included in the experimental protocol to ensure that bacteria of the low inoculum model had time to revert to log-phase growth from their original early plateau phase source culture. However, this made accurately controlling the initial cfu/mL values in the HFIM in each group challenging, as the cells of the two cultures exhibited different doubling times and subsequently had unequal growth over the acclimatisation period. This led to lower than intended cfu/mL value separation between the initial high and low inoculum groups in some experiments; most notably the populations receiving bolus-dosed benzylpenicillin. The separation deficit had the potential to mask or reduce any difference in the two groups. However, the obtained results showed significant differences in antibiotic susceptibility between the groups at 24 hours, suggesting that growth phase of the bacterial population has greater effect on the manifestation of the inoculum effect than absolute cell density. It is uncertain whether this conclusion can be extended to other antibiotics – it is possible that in some antibiotics the advent of the effect may be growth phase-dependent (as it appears to be with penicillin), while cell count may be of primary importance in others.

### Use of Cation-adjusted Mueller-Hinton Broth

Strep A has been previously thought to be only marginally susceptible to trimethoprim-sulfamethoxazole. This may have been due to the media on which its susceptibility has historically been assayed. SXT acts by blocking two steps in folic acid metabolism, preventing downstream thymidine synthesis, a mechanism which may have been circumvented by performing susceptibility testing on media containing thymidine. Cation-adjusted Mueller-Hinton broth was chosen for its low, defined thymidine content which would not mask susceptibility, and under these conditions the results showed that Strep A is susceptible to SXT. It is unknown whether this observation can be generalised to *in vivo* susceptibility, as serum thymidine concentration cannot be regulated to the extent possible in artificial media. However, previous studies have measured the human serum concentration of thymidine at approximately 0.02-0.01 μg/mL^25^, lower than that present in Mueller-Hinton blood agar (MHBA), which when cultured with Strep A produces SXT-susceptible organisms^12^.

### Validation of the HFIM of *Strep A* Infection

Previous experimental data on Strep A infection and data produced in the hollow fibre infection model during this research is mostly in agreement. An inoculum effect has previously been reported^5^ in penicillin in a murine model of Strep A myositis, as well as being seen clinically as a cause of penicillin monotherapy failure. Our results support this conclusion, with the continuously infused benzylpenicillin group displaying an inoculum effect at 24 and 48 hours, and in the bolus-dosed group at 24 hours. As previously highlighted, the effect was not seen at 48 hours, possibly due to the loss of quorum sensing and other signalling molecules by the high flow of media, which prevented the advent of the inoculum effect after bacterial population fell below a certain level. As such, the use of the HFIM for Strep A research may be better suited to antibiotics with lower clearance rates, or high clearance rate antibiotics alongside high bacterial concentrations.

The generalisability of the HFIM to clinical research was further borne out by the results of the clindamycin experiment, which showed no inoculum effect at 24 hours and one of small magnitude at 48 hours. Previous literature has been mixed in reporting an inoculum effect, with *in vitro* studies^6^ showing an effect (albeit generally of lesser magnitude than seen in benzylpenicillin), while murine models have failed to demonstrate this.^5^ These results suggest that the HFIM may represent a more nuanced approximation of Strep A infection that is closer to an *in vivo* model than previously achieved *in vitro*.

Our literature search was unable to identify any studies on the inoculum effect of linezolid in Strep A. However, murine models of *Streptococcus pneumoniae* and *Staphylococcus aureus* treated with linezolid have noted the effect.^23^ Further research is needed to confirm whether it is present in *in vivo* models of Strep A infection as is suggested by the HFIM model. Similarly, no animal studies trialling SXT for an inoculum effect were found, although a modest effect was seen for both Strep A and *Haemophilus influenzae* in *in vitro* testing.^24^ This is a possible point of discrepancy between the HFIM and previous literature; additional investigation must be completed in animal models to assess the relevance of the HFIM to the study of SXT inoculum effects *in vivo*. Simulation of combination therapy should also be explored and would be feasible with the HFIM.

## Acknowledgements

This work was supported by the Spinnaker Health Research Foundation (FHMRF 2016).

## Transparency and Declarations

None to declare

## Footnotes

* Sample meta-data are available in the BioSamples database (http://www.ebi.ac.uk/biosamples) under accession number SAMEA1523579, or at https://www.ebi.ac.uk/biosamples/samples/SAMEA1523579

